# HIF-1α/STOML2 mediated PINK1-dependent mitophagy activation against hypoxia-induced neuronal injury

**DOI:** 10.1101/2025.04.16.649120

**Authors:** Yuning Li, Mengyuan Guo, Zhengming Tian, Qianqian Shao, Yingxia Liu, Yakun Gu, Zirui Xu, Feiyang Jin, Xunming Ji, Jia Liu

**Affiliations:** Beijing Institute of Brain Disorders, Laboratory of Brain Disorders, Hypoxia Conditioning Translational Laboratory of Clinical Medicine, Ministry of Science and Technology, Collaborative Innovation Center for Brain Disorders, Beijing Advanced Innovation Center for Big Data-based Precision Medicine, Capital Medical University, Beijing, China; Department of Neurosurgery, Xuanwu Hospital, Capital Medical University, Beijing, China

**Author notes:** Corresponding author: Jia Liu,.

**Keywords:** Hypoxia, Mitophagy, PINK1, HIF-1α, Intermittent hypoxia

## Abstract

Hypoxia contributes to brain disorders by causing neuronal injury. However, in the early stage of stress, neurons initiate a series of compensatory pathways to resist cell damage, but the underlying mechanisms have not been fully elucidated. In this study, we found that hypoxia transiently activates PTEN-induced kinase 1 (PINK1)-dependent mitophagy in the early stage before cell damage and neurological dysfunction. PINK1 overexpression protects neurons and it knockdown exacerbates neuronal damage, highlighting the key role of PINK1-dependent mitophagy in hypoxic adaptation. Mechanistically, hypoxia promotes HIF-1α nuclear translocation, inducing transcription of stomatin like 2 (STOML2). STOML2 relocates to the mitochondrial membrane, aiding phosphoglycerate mutase 5 (PGAM5) cleavage, which triggers PINK1-dependent mitophagy. Silencing HIF-1α, STOML2, or PGAM5 inhibits PINK1-dependent mitophagy and worsens neurological function under hypoxia. Notably, intermittent hypoxia, a hypoxic conditioning strategy for improving hypoxic tolerance, enhances PINK1-dependent mitophagy by activating HIF-1α/STOML2 axis, and protects neurons against hypoxia. In conclusion, our study reveals a new “self-protection” mechanism of neurons against hypoxic stress and discovers that intermittent hypoxia is a potential therapeutic strategy against hypoxia-induced injury.

## Introduction

Hypoxia is implicated in a variety of central nervous system (CNS) disorders, such as stroke and Parkinson’s disease, suggesting that hypoxia may act as a co-factor in neurological injury(Li *et al*, 2017). One of the primary mechanisms by which hypoxia induces damage is through the disruption of mitochondrial function(Li *et al*, 2025). Mitochondria, the cell’s energy powerhouse, are responsible for oxidative phosphorylation, the process that produces ATP (adenosine triphosphate), the cell’s main energy currency(Alshial *et al*, 2023). The efficiency of this process is critical for maintaining cellular homeostasis, particularly in energy-demanding tissues like the brain(Hoffmann *et al*, 2021; Schmitt & Gaspar, 2023). Under hypoxic conditions, mitochondrial dysfunction is commonly observed and has been implicated as a central pathogenic factor in several neurological disorders(Liang *et al*, 2024). Dysfunctional mitochondria lead to the accumulation of reactive oxygen species (ROS), changes in mitochondrial dynamics, and eventually, cellular apoptosis(Almeida, 2024; Wen *et al*, 2025). The accumulation of these damaged mitochondria contributes to neuronal injury in hypoxic environments. Therefore, understanding how hypoxia disrupts mitochondrial function and exploring the cellular mechanisms that mitigate this damage could reveal novel therapeutic targets for hypoxia-related neurological injuries.

One critical mechanism for mitigating mitochondrial dysfunction is mitophagy, a selective form of autophagy that plays a crucial role in maintaining mitochondrial quality control by targeting and degrading dysfunctional or damaged mitochondria(Szczepanowska & Trifunovic, 2022). Although macroautophagy has similar functions, mitophagy is more specific and effective in dealing with dysfunctional mitochondria(Tian *et al*, 2025). Mitophagy is essential for the proper maintenance of cellular homeostasis, particularly in high-energy-demanding tissues such as neurons and muscles, where the proper functioning of mitochondria is critical for survival(Cheng *et al*, 2025; Meng *et al*, 2025). Currently, a literature review summarizes the factors affecting mitophagy, among which hypoxia is an important way to activate mitophagy(Wu & Chen, 2015). Studies have shown that FUNDC1 (FUN14 domain-containing protein 1) is the classical mitophagy receptor in response to hypoxia(Kuang *et al*, 2016; Liu *et al*, 2012). However, after conducting more literature searches, we found that not only FUNDC1 but also a variety of other factors can mediate the activation of mitophagy under hypoxic stress. For example, PINK1 (PTEN-induced kinase 1), PINK1 is a classical mitophagy receptor(Ge *et al*, 2020). Hypoxia has been shown to promote PINK1-dependent mitophagy(Linqing *et al*, 2021). Although it is known that hypoxia promotes PINK1 accumulation and mitophagy activation, the exact mechanisms remain unclear. Understanding these mechanisms could lead to the identification of novel therapeutic targets for treating diseases related to mitochondrial dysfunction under hypoxic stress.

In this study, we found that early-stage hypoxia activates PINK1-dependent mitophagy through the HIF-1α/STOML2/PGAM5 pathway, providing neuroprotection against hypoxic damage. To further investigate this, we performed knockdown experiments targeting these molecules. Additionally, we observed that intermittent hypoxia (IH) may also exert neuroprotective effects by activating this mitophagy pathway, offering promising clinical applications in the treatment of neurodegenerative diseases and conditions involving mitochondrial dysfunction.

## Results

### Mitophagy undergoes transient compensatory activation in the early hypoxic stage to resist cell damage

Adult C57BL mice were continuously exposed to 13% O₂, the oxygen concentration found on the Tibetan Plateau, for 0, 1, 3, and 7 days (Con, H1d, H3d, H7d). And we identify the H1d and H3d as early stages of hypoxia, while H7d as long-term hypoxic treatment. They subsequently underwent behavioral and postmortem histological analyses to assess neurological damage (Figure 1A). To evaluate cognitive function, we performed the Novel Object Test, which evaluate novel object exploration and spatial exploration, respectively. Behavioral analysis revealed that while H1d and H3d mice maintained normal cognitive performance, H7d mice exhibited significant cognitive decline, indicating that prolonged hypoxia impairs cognitive function (Figure 1B–C). Histological analysis of hippocampal neurons using Nissl staining revealed that neuronal arrangements in the Con, H1d, and H3d groups were orderly, whereas H7d mice exhibited disorganized hippocampal neuronal structures (Figure 1D). Quantification of Nissl-positive cells, which represent viable neurons, further confirmed a significant decrease exclusively in the H7d group (Figure 1E). These findings align with previous reports of cognitive impairment associated with prolonged hypoxia, suggesting that hypoxia-induced neuronal damage does not occur immediately. Instead, certain protective mechanisms may allow neurons to withstand early-stage hypoxic conditions. To investigate whether mitophagy contributes to hypoxia resistance, we first assessed LC3-I and LC3-II levels in mitochondrial fractions via western blot analysis (Figure 1F). The conversion of LC3-I to LC3-II serves as a marker of mitophagy activation. An increase in LC3-II/I ratio indicates mitophagy activation, which was significantly elevated in H1d and H3d groups (Figure 1G). This suggests that mitophagy may play a crucial role in protecting against hypoxia-induced cognitive impairment. In summary, mitophagy is activated during the early stages of hypoxia (1–3 days) and may play a critical role in protecting against hypoxia-induced cognitive impairment.

**Figure 1.**
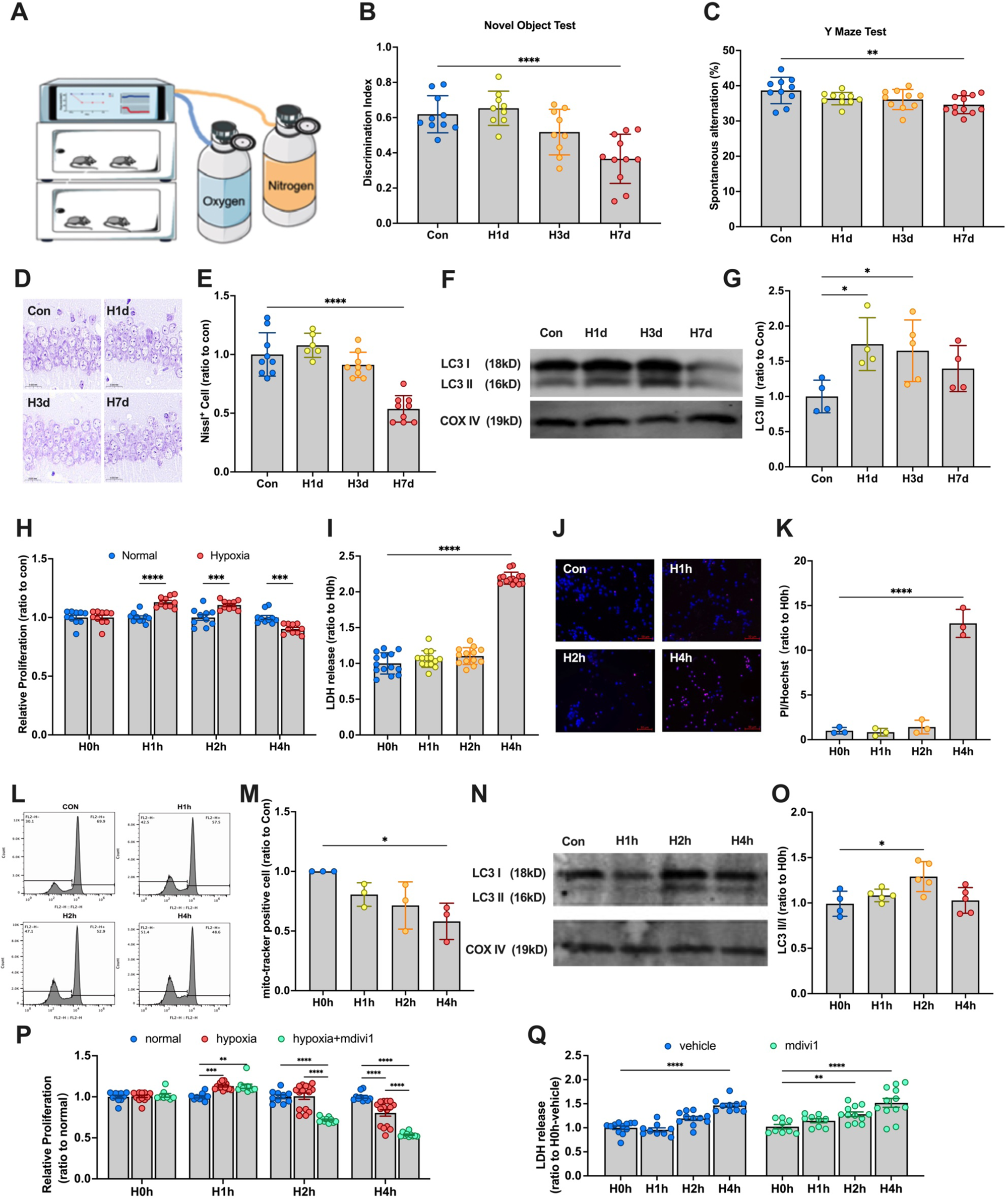
Neurons resist cellular damage by activating mitophagy in the early stage of hypoxic. **(A)** Specific hypoxia patterns. Mice were placed in hypoxic chambers with an O_2_ concentration of 13%. Constant levels of nitrogen and oxygen were maintained to achieve persistent hypoxia. **(B-C)** Mice were continuously treated with hypoxia for 0,1, 3, and 7 days (Con, H1d, H3d, H7d), and behavioral tests were performed at different time points. **(B)** Detection and statistical analysis of Novel Object Test in mice of each group. **(C)** Detection and statistical analysis of the Y Maze Test in mice of each group. **(D)** Nissl staining was used to evaluate the neuronal arrangement in the hippocampus. **(E)** The levels of Nissl-positive cells in mice of each group. **(F)** The levels of LC3 in the hippocampus mitochondrial protein were detected by western blotting using COX IV as the internal reference. **(G)** Statistical analysis of LC3 II/I in mice of each group. Statistical analysis of western blot was performed by homogenization, which means setting the value of the Con group as 1. **(H)** SY5Y cells were cultured with 21% O_2_(normal) and 1% O_2_ (hypoxia) for 0, 1, 2 and 4h, the cell relative proliferation was detected using the CCK-8 assay. **(I)** SY5Y cells were cultured with 1% O_2_ for 0, 1, 2 and 4h, the cytotoxicity was detected using LDH assay. **(J)** The effect of hypoxic treatment on apoptosis was observed using PI/Hoechst staining kit and observed under a fluorescence microscope. **(K)** Statistical analysis of PI/Hoechst in different groups. **(L)** After mito-tracker staining, detected the mitochondria damage in SY5Y cells by flow cytometry. **(M)** Statistical analysis of healthy mitochondria counting. **(N)** The levels of LC3 in the mitochondrial protein in SY5Y cells were detected by western blotting using COX IV as the internal reference. **(O)** Statistical analysis of LC3 II/I in mice of each group. Statistical analysis of western blot was performed by homogenization, which means setting the value of the H0h group as 1. **(P)** The cell relative proliferation was detected using the CCK-8 assay after suppressing mitophagy by mdivi1. Analysis was performed by homogenization, which means setting the value of the normal group as 1. **(Q)** The cytotoxicity was detected using LDH assay after suppressing mitophagy by mdivi1. Data was analyzed via one-way ANOVA and Tukey’s post-test. *p < 0.05, **p < 0.01, ***p < 0.001, ****p < 0.0001.

To evaluate neuronal responses to hypoxia, we utilized the SH-SY5Y cell line and cultured the cells in 1% O₂. Cell viability was assessed using the CCK-8 assay, which revealed a significant increase in proliferation in the H1h and H2h groups, whereas a significant decline was observed in the H4h group (Figure 1H). Cytotoxicity was assessed using the LDH assay, which showed a significant increase in LDH release exclusively in the H4h group (Figure 1I). Apoptosis rates were evaluated using PI/Hoechst staining, with analysis revealing a marked increase in the PI/Hoechst ratio in the H4h group (Figure 1J–K). These results are consistent with findings in mice, further supporting that hypoxia-induced neuronal damage occurs at a later phase of hypoxic exposure. To assess mitochondrial integrity, cells were stained with mito-tracker dye and analyzed by flow cytometry. The number of mito-tracker-positive (healthy) mitochondria was significantly reduced in the H4h group, indicating mitochondrial damage. To determine whether mitophagy protects against hypoxia-induced damage, we assessed LC3-II/I levels via western blot analysis (Figure 1N) and found a significant increase in LC3-II/I in the H2h group (Figure 1O). Furthermore, to confirm the functional role of mitophagy, we inhibited mitophagy using mdivi-1. After mdivi-1 treatment, cell viability significantly decreased in the H2h group, suggesting that blocking mitophagy accelerates hypoxia-induced neuronal damage. Moreover, LDH release was significantly increased in H2h as well as H4h, further indicating that mitophagy is critical for neuronal survival under hypoxia. In summary, mitophagy is activated in neurons under hypoxic conditions and plays a protective role in preventing hypoxia-induced damage.

### PINK1-dependent mitophagy is an indispensable pathway for neurons to resist cell damage induced by hypoxia

To identify key mediators of hypoxia-induced mitophagy, we examined whether PINK1 (PTEN-induced kinase 1) regulates mitophagy activation. As a classical mitophagy receptor (Ge *et al*., 2020), PINK1 has been implicated in hypoxia-induced mitophagy, though the precise activation mechanism remains unclear. To address this, we isolated mitochondrial fractions and analyzed PINK1 levels. Western blot analysis revealed a significant increase in mitochondrial PINK1 expression in H1d and H3d mice (Figure 2A–B) and H2h cells (Figure 2C–D), confirming hypoxia-induced PINK1 upregulation. These findings suggest that PINK1-dependent mitophagy plays a crucial role in hypoxic adaptation.

**Figure 2.**
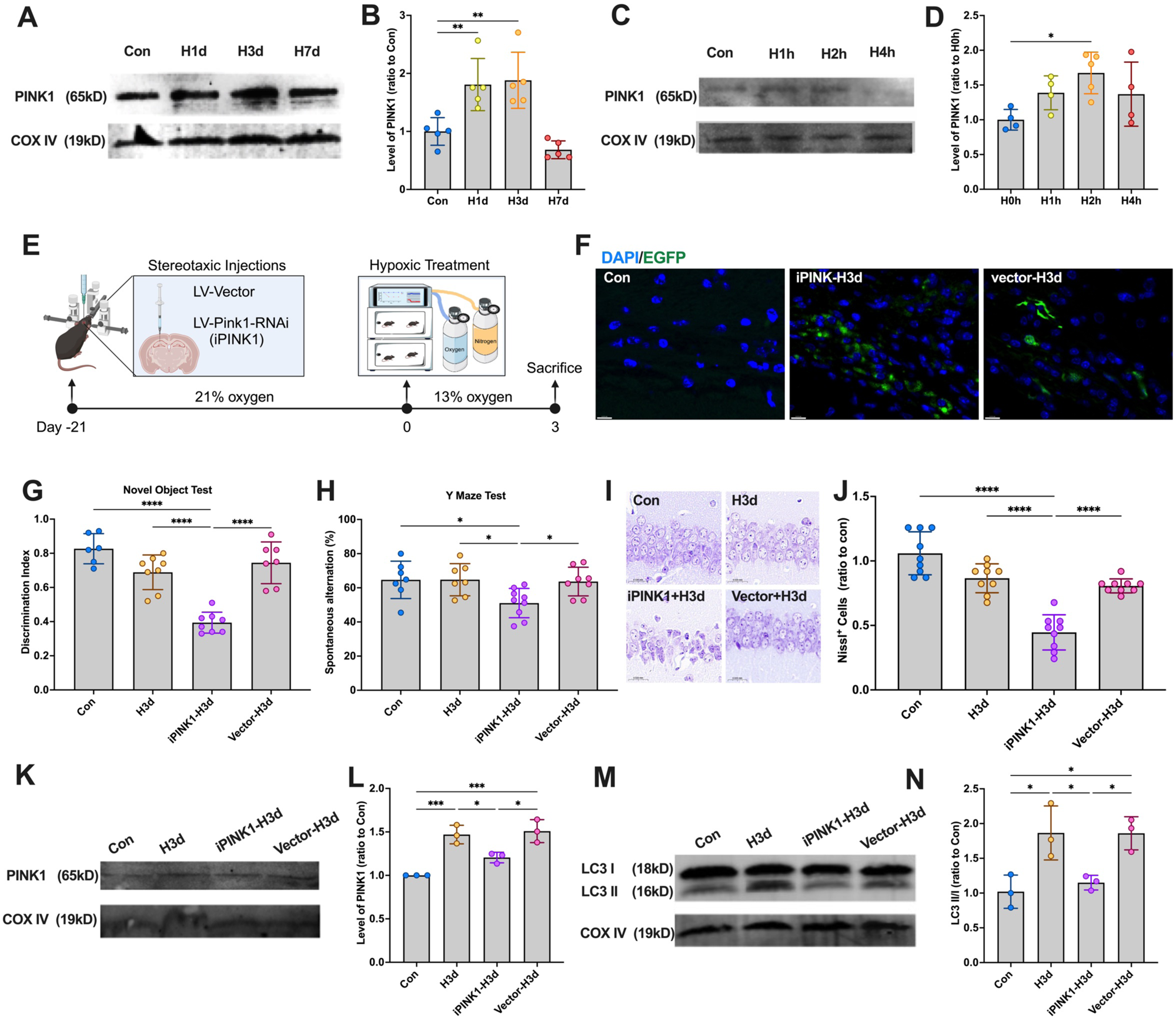
PINK1-dependent mitophagy is a necessary pathway helping neurons to resist hypoxic damage. **(A)** Mice were continuously treated with hypoxia for 0,1, 3, and 7 days (Con, H1d, H3d, H7d), and the levels of PINK1 in the hippocampus mitochondrial protein were detected by western blotting using COX IV as the internal reference. **(B)** Statistical analysis of PINK1 in mice of each group. Statistical analysis of western blot was performed by homogenization, which means setting the value of the Con group as 1. **(C)** SY5Y cells were continuously treated with hypoxia for 0,1, 2, and 4 hours (H0h, H1h, H2h, H4h), and the levels of PINK1 in the neuron mitochondrial protein were detected by western blotting using COX IV as the internal reference. **(D)** Statistical analysis of PINK1 in cells of each group. Statistical analysis of western blot was performed by homogenization, which means setting the value of the H0h group as 1. **(E-N)** Mice were administered the lentiviral targeting PINK1 and then received continuous hypoxic treatment for 3 days. Mice were grouped in 4 (Con, H3d, iPINK1-H3d and vector-H3d). **(E)** The timeline of lentiviral injection and hypoxic treatment. **(F)** Expression of LV-Pink1-RNAi-EGFP in the hippocampus was observed under a fluorescence microscope. **(G)** Detection and statistical analysis of Novel Object Test in mice of each group. **(H)** Detection and statistical analysis of the Y Maze Test in mice of each group. **(I)** Nissl staining was used to evaluate the neuronal arrangement in the hippocampus. **(J)** The levels of Nissl-positive cells in mice of each group. **(K)** The levels of PINK1 in mice of each group in the hippocampus mitochondrial protein were detected by western blotting using COX IV as the internal reference. **(L)** Statistical analysis of PINK1 in mice of each group. Statistical analysis of western blot was performed by homogenization, which means setting the value of the Con group as 1. **(M)** The levels of LC3 in the hippocampus mitochondrial protein were detected by western blotting using COX IV as the internal reference. **(N)** Statistical analysis of LC3 II/I in mice of each group. Data was analyzed via one-way ANOVA and Tukey’s post-test. *p < 0.05, **p < 0.01, ***p < 0.001, ****p < 0.0001.

To validate PINK1’s role, mice were administered lentiviral constructs targeting PINK1 and exposed to hypoxia for 3 days before undergoing behavioral and histological analysis (Figure 2E). EGFP labeling confirmed successful viral transduction (Figure 2F). As expected, iPINK1-H3d mice exhibited cognitive decline, while H3d and vector-H3d mice showed no significant impairment (Figure 2G–H). Consistently, Nissl staining revealed significant neuronal damage in iPINK1-H3d mice, whereas H3d and vector-H3d groups maintained hippocampal integrity (Figure 2I–J). These findings indicate that PINK1 protects against hypoxia-induced neuronal damage.

To confirm the role of PINK1 in mitophagy activation, we performed a western blot analysis to assess its expression in mitochondrial fractions (Figure 2K–L). The results showed that PINK1 was significantly upregulated in H3d but was reduced upon inhibition. Next, to determine whether PINK1 mediates hypoxia-induced mitophagy, we analyzed LC3 expression via western blot (Figure 2M). Results demonstrated that mitophagy activation, indicated by increased LC3-II/I ratios, was abolished in iPINK1-H3d mice, confirming that hypoxia-induced mitophagy is PINK1-dependent (Figure 2N). Meanwhile, after injection of PINK1 overexpressing lentivirus, we found that the increase in mitophagy level would not be inhibited (Figure S1), further suggesting that early hypoxia did activate PINK1-dependent mitophagy.

### Full-length PGAM5 mediates PINK1-dependent mitophagy under hypoxia

Early hypoxia activates PINK1-dependent mitophagy to mitigate neuronal injury. However, the mechanism by which PINK1 is stabilized on the outer mitochondrial membrane (OMM) and its functional role remains unclear.

Our early research showed that PGAM5 and STOML2 significantly increased after Hypoxia. These two factors are associated with mitophagy(Shao *et al*, 2022). PGAM5 is a serine/threon ine phosphatase on mitochondria(Cheng *et al*, 2021), and previous studies suggest that full-length PGAM5 (L-PGAM5) facilitates PINK1 stabilization on the OMM(Yan *et al*, 2020; Zeb *et al*, 2021). To explore the role of L-PGAM5 in early hypoxia-induced mitophagy, we isolated mitochondrial proteins from hippocampal tissues and SH-SY5Y cells for western blot analysis. Results showed a significant increase in mitochondrial L-PGAM5 levels after 3 days of hypoxia in mice (Figure 3A-B) and after 1–2 hours of hypoxia in cells (Figure 3C-D).

**Figure 3.**
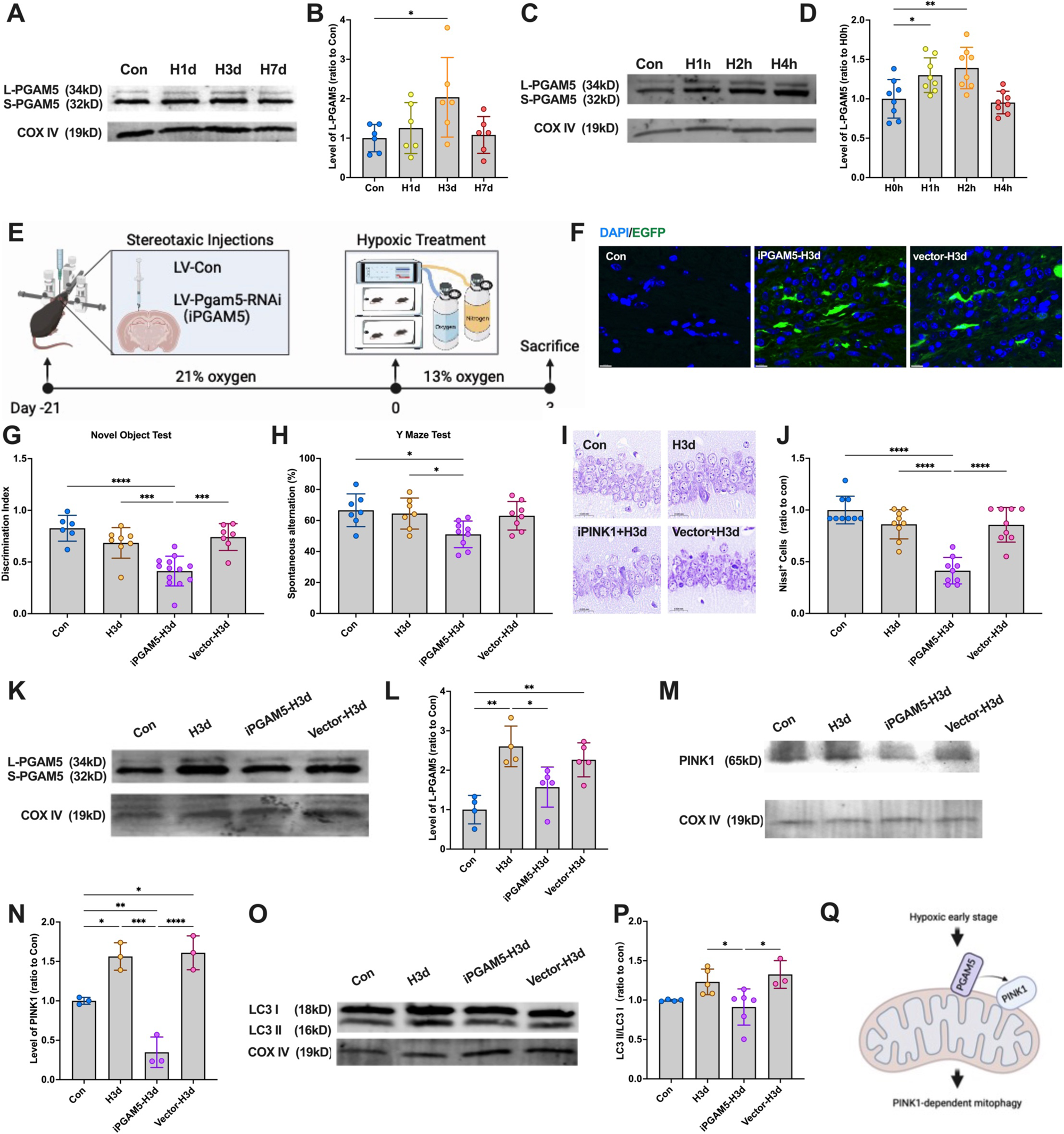
L-PGAM5 mediates PINK1-dependent mitophagy. **(A)** Mice were continuously treated with hypoxia for 0,1, 3, and 7 days (Con, H1d, H3d, H7d), and the levels of PGAM5 in the hippocampus mitochondrial protein were detected by western blotting using COX IV as the internal reference. **(B)** Statistical analysis of L-PGAM5 in mice of each group. Statistical analysis of western blot was performed by homogenization, which means setting the value of the Con group as 1. **(C)** SY5Y cells were continuously treated with hypoxia for 0,1, 2, and 4 hours (H0h, H1h, H2h, H4h), and the levels of PGAM5 in the neuron mitochondrial protein were detected by western blotting using COX IV as the internal reference. **(D)** Statistical analysis of L-PGAM5 in cells of each group. Statistical analysis of western blot was performed by homogenization, which means setting the value of the H0h group as 1. **(E-N)** Mice were administered the lentiviral targeting PGAM5 and then received continuous hypoxic treatment for 3 days. Mice were grouped in 4 (Con, H3d, iPGAM5-H3d and vector-H3d). **(E)** The timeline of lentiviral injection and hypoxic treatment. **(F)** Expression of LV-Pgam5-RNAi-EGFP in the hippocampus was observed under a fluorescence microscope. **(G)** Detection and statistical analysis of Novel Object Test in mice of each group. **(H)** Detection and statistical analysis of the Y Maze Test in mice of each group. **(I)** Nissl staining was used to evaluate the neuronal arrangement in the hippocampus. **(J)** The levels of Nissl-positive cells in mice of each group. **(K)** The levels of L-PGAM5 in mice of each group in the hippocampus mitochondrial protein were detected by western blotting using COX IV as the internal reference. **(L)** Statistical analysis of L-PGAM5 in mice of each group. **(M)** The levels of PINK1 in mice of each group in the hippocampus mitochondrial protein were detected by western blotting using COX IV as the internal reference. **(N)** Statistical analysis of PINK1 in mice of each group. **(O)** The levels of LC3 in the hippocampus mitochondrial protein were detected by western blotting using COX IV as the internal reference. **(P)** Statistical analysis of LC3 II/I in mice of each group. **(Q)** The mechanism diagram of the pathway. Data was analyzed via one-way ANOVA and Tukey’s post-test. *p < 0.05, **p < 0.01, ***p < 0.001, ****p < 0.0001.

To determine whether PINK1 residency on mitochondria is necessary for mitophagy activation, we knocked down PGAM5 in the hippocampus via lentiviral injection (LV-Pgam5-RNAi) using stereotactic surgery and allowed three weeks for viral expression before hypoxia exposure (Figure 3E). EGFP labeling confirmed successful viral transduction (Figure 3F). As expected, iPGAM5-H3d mice exhibited cognitive decline, whereas H3d and vector-H3d mice showed no significant impairment (Figure 3G–H). Nissl staining further revealed significant neuronal damage in iPGAM5-H3d mice, while hippocampal integrity was preserved in H3d and vector-H3d groups (Figure 3I–J). These findings suggest that PGAM5 protects against hypoxia-induced neuronal damage.

Next, we performed western blot analysis of mitochondrial proteins to confirm PGAM5 knockdown efficiency (Figure 3K). L-PGAM5 levels were significantly reduced in the iPGAM5-H3d group, confirming successful knockdown (Figure 3L). To investigate whether L-PGAM5 acts upstream of PINK1-dependent mitophagy, we examined mitochondrial PINK1 and LC3 levels in PGAM5-knockdown mice. PGAM5 depletion reversed the hypoxia-induced increase in mitochondrial PINK1 (Figure 3M-N) and abolished mitophagy activation (Figure 3O-P) in the early stages of hypoxia. These findings indicate that L-PGAM5 upregulation during early hypoxia is crucial for PINK1 stabilization and mitophagy activation (Figure 3Q).

### Increased STOML2 on mitochondria upregulates full-length PGAM5 level

Our experimental results confirm that early hypoxia increases mitochondrial L-PGAM5, which activates PINK1-dependent mitophagy to protect against hypoxic injury. However, the mechanism by which L-PGAM5 is stabilized on mitochondria remains unclear. Previous studies suggest that STOML2, which increased significantly after hypoxia, is widely recognized for its role in mitochondrial biogenesis and inner membrane organization during cancer growing(Christie *et al*, 2012; Fan *et al*, 2022; Guo *et al*, 2022; Ma *et al*, 2021), such as gastric cancer. However, the role of STOML2 in the nervous system is currently unknown. STOML2 has been implicated in PGAM5 cleavage regulation (Wai *et al*, 2016), suggesting a potential upstream role in mitophagy activation.

To investigate STOML2’s role, we examined its mitochondrial levels under early hypoxia using western blot. Results showed a significant increase in mitochondrial STOML2 in H3d mice (Figure 4A-B) and in SH-SY5Y cells after 2 hours of hypoxia (H2h) (Figure 4C-D), suggesting that STOML2 upregulation accompanies L-PGAM5-mediated activation of PINK1-dependent mitophagy.

**Figure 4.**
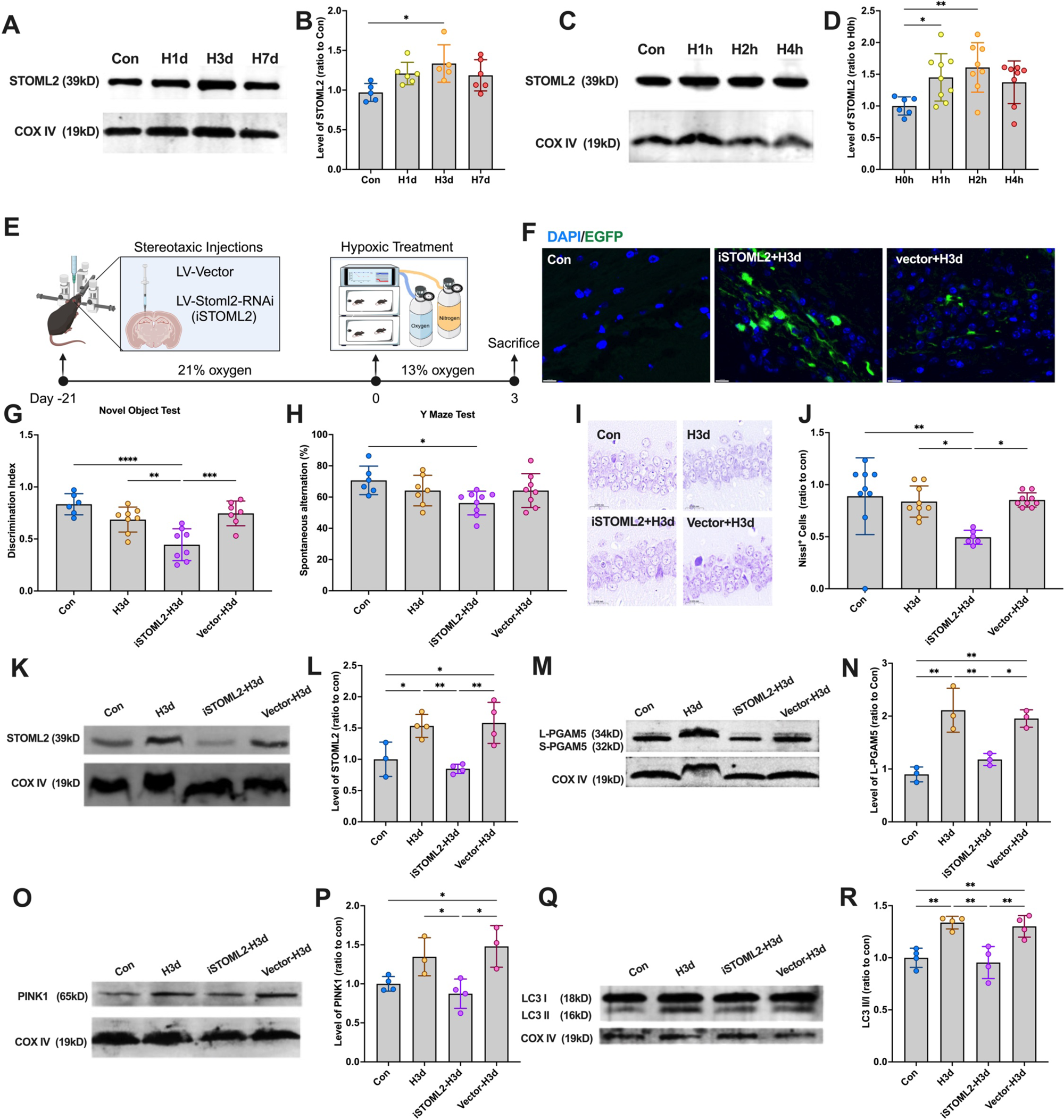
STOML2 helps L-PGAM5 to be stable on the membrane of mitochondria. **(A)** Mice were continuously treated with hypoxia for 0,1, 3, and 7 days (Con, H1d, H3d, H7d), and the levels of STOML2 in the hippocampus mitochondrial protein were detected by western blotting using COX IV as the internal reference. **(B)** Statistical analysis of STOML2 in mice of each group. Statistical analysis of western blot was performed by homogenization, which means setting the value of the Con group as 1. **(C)** SY5Y cells were continuously treated with hypoxia for 0,1, 2, and 4 hours (H0h, H1h, H2h, H4h), and the levels of STOML2 in the neuron mitochondrial protein were detected by western blotting using COX IV as the internal reference. **(D)** Statistical analysis of STOML2 in cells of each group. Statistical analysis of western blot was performed by homogenization, which means setting the value of the H0h group as 1. **(E-N)** Mice were administered the lentiviral targeting STOML2 and then received continuous hypoxic treatment for 3 days. Mice were grouped in 4 (Con, H3d, iSTOML2-H3d and vector-H3d). **(E)** The timeline of lentiviral injection and hypoxic treatment. **(F)** Expression of LV-Stoml2-RNAi-EGFP in the hippocampus was observed under a fluorescence microscope. **(G)** Detection and statistical analysis of Novel Object Test in mice of each group. **(H)** Detection and statistical analysis of the Y Maze Test in mice of each group. **(I)** Nissl staining was used to evaluate the neuronal arrangement in the hippocampus. **(J)** The levels of Nissl-positive cells in mice of each group. **(K)** The levels of STOML2 in mice of each group in the hippocampus mitochondrial protein were detected by western blotting using COX IV as the internal reference. **(L)** Statistical analysis of STOML2 in mice of each group. **(M)** The levels of L-PGAM5 in mice of each group in the hippocampus mitochondrial protein were detected by western blotting using COX IV as the internal reference. **(N)** Statistical analysis of L-PGAM5 in mice of each group. **(O)** The levels of PINK1 in mice of each group in the hippocampus mitochondrial protein were detected by western blotting using COX IV as the internal reference. **(P)** Statistical analysis of PINK1 in mice of each group. **(Q)** The levels of LC3 in the hippocampus mitochondrial protein were detected by western blotting using COX IV as the internal reference. **(R)** Statistical analysis of LC3 II/I in mice of each group. Data was analyzed via one-way ANOVA and Tukey’s post-test. *p < 0.05, **p < 0.01, ***p < 0.001, ****p < 0.0001.

To determine whether STOML2 stabilizes L-PGAM5 on mitochondria, we knocked down STOML2 in the hippocampus (LV-Stoml2-RNAi) via stereotactic injection and allowed three weeks for viral expression before hypoxia exposure (Figure 4E). EGFP labeling confirmed successful transduction (Figure 4F). As expected, iSTOML2-H3d mice exhibited cognitive decline, whereas H3d and vector-H3d mice showed no significant impairment (Figure 4G–H). Nissl staining further revealed significant neuronal damage in iSTOML2-H3d mice, while hippocampal integrity was maintained in H3d and vector-H3d groups (Figure 4I–J). These findings suggest that STOML2 protects against hypoxia-induced neuronal damage.

To confirm the functional role of STOML2 in mitophagy regulation, we knocked down STOML2 and assessed its mitochondrial protein levels via western blot (Figure 4K–L). Western blot analysis confirmed that STOML2 was successfully depleted in iSTOML2-H3d mice. Next, we examined whether STOML2 is necessary for L-PGAM5 stabilization by analyzing mitochondrial protein fractions (Figure 4M–N). Meanwhile, our results showed that the knockdown of PGAM5 did not affect STOML2 expression (Figure S2). The results showed that STOML2 knockdown reversed the hypoxia-induced increase in L-PGAM5 expression, indicating that STOML2 functions upstream of PGAM5 in mitophagy activation.

Next, we examined whether STOML2 upregulation enhances PINK1-dependent mitophagy. Western blot analysis revealed that STOML2 knockdown abolished hypoxia-induced PINK1 accumulation on mitochondria (Figure 4O-P) and prevented mitophagy activation (Figure 4Q-R). Together, these findings demonstrate that hypoxia-induced STOML2 upregulation stabilizes L-PGAM5, which in turn activates PINK1-dependent mitophagy to confer resistance against hypoxic injury. STOML2 is widely recognized for its role in mitochondrial biogenesis and inner membrane organization during cancer growing[21, 22]. However, our study uncovers a previously unknown function of STOML2 in regulating mitophagy under hypoxia.

### Hypoxia promotes STOML2 transcription through HIF-1α pathway and thus activates mitophagy

To investigate HIF-1α nuclear translocation under hypoxia, we analyzed HIF-1α levels in nuclear fractions from mouse hippocampal tissues and SH-SY5Y cells using western blot. Results showed a significant increase in nuclear HIF-1α in H1d and H3d mice (Figure 5A-B) and in H2h cells (Figure 5C-D), indicating that early hypoxia promotes HIF-1α nuclear entry. This suggests that hypoxia-induced mitophagy may be HIF-1α-dependent.

**Figure 5.**
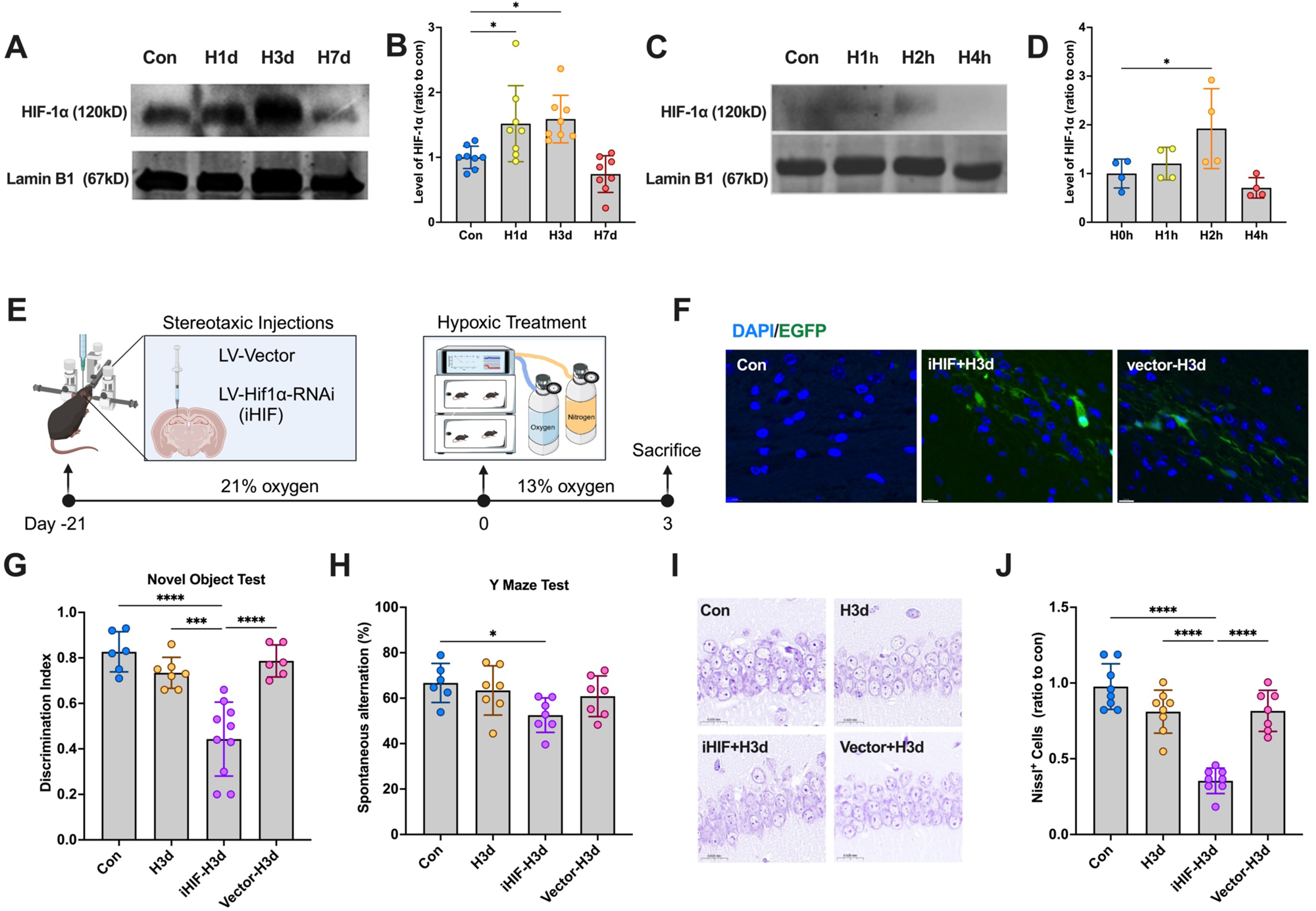
Neurons resist cellular damage by activating HIF-1α. **(A)** Mice were continuously treated with hypoxia for 0,1, 3, and 7 days (Con, H1d, H3d, H7d), and the levels of HIF1α in the hippocampus nucleus protein were detected by western blotting using LaminB1 as the internal reference. **(B)** Statistical analysis of HIF1α in mice of each group. Statistical analysis of western blot was performed by homogenization, which means setting the value of the Con group as 1. **(C)** SY5Y cells were continuously treated with hypoxia for 0,1, 2, and 4 hours (H0h, H1h, H2h, H4h), and the levels of HIF1α in the neuron mitochondrial protein were detected by western blotting using COX IV as the internal reference. **(D)** Statistical analysis of HIF1α in cells of each group. Statistical analysis of western blot was performed by homogenization, which means setting the value of the H0h group as 1. **(E-N)** Mice were administered the lentiviral targeting HIF1α and then received continuous hypoxic treatment for 3 days. Mice were grouped in 4 (Con, H3d, iHIF1α-H3d and vector-H3d). **(E)** The timeline of lentiviral injection and hypoxic treatment. **(F)** Expression of LV-Hif1α-RNAi-EGFP in the hippocampus was observed under a fluorescence microscope. **(G)** Detection and statistical analysis of Novel Object Test in mice of each group. **(H)** Detection and statistical analysis of the Y Maze Test in mice of each group. **(I)** Nissl staining was used to evaluate the neuronal arrangement in the hippocampus. **(J)** The levels of Nissl-positive cells in mice of each group. Data was analyzed via one-way ANOVA and Tukey’s post-test. *p < 0.05, **p < 0.01, ***p < 0.001, ****p < 0.0001.

To confirm HIF-1α’s role, we knocked down HIF-1α in the hippocampus (LV-Hif1α-RNAi) via stereotactic injection and allowed three weeks for viral expression before hypoxia exposure (Figure 5E). EGFP labeling confirmed successful transduction (Figure 5F). As expected, iHIF-H3d mice exhibited cognitive decline, while H3d and vector-H3d mice showed no significant impairment (Figure 5G–H). Nissl staining further revealed significant neuronal damage in iHIF-H3d mice, whereas hippocampal integrity was maintained in H3d and vector-H3d groups (Figure 5I–J). These findings indicate that HIF-1α protects against hypoxia-induced neuronal damage.

STOML2 has been identified as a novel downstream target of HIF-1α [5]. To assess STOML2 transcriptional regulation by HIF-1α, we measured STOML2 mRNA levels via qPCR. In both mice (H3d) and SH-SY5Y cells (H30min, H60min, H90min), STOML2 mRNA was significantly increased, peaking at 60 minutes of hypoxia (Figure 6A-B). Notably, HIF-1α knockdown abolished this upregulation (Figure 6C), confirming that HIF-1α nuclear translocation enhances STOML2 transcription.

**Figure 6.**
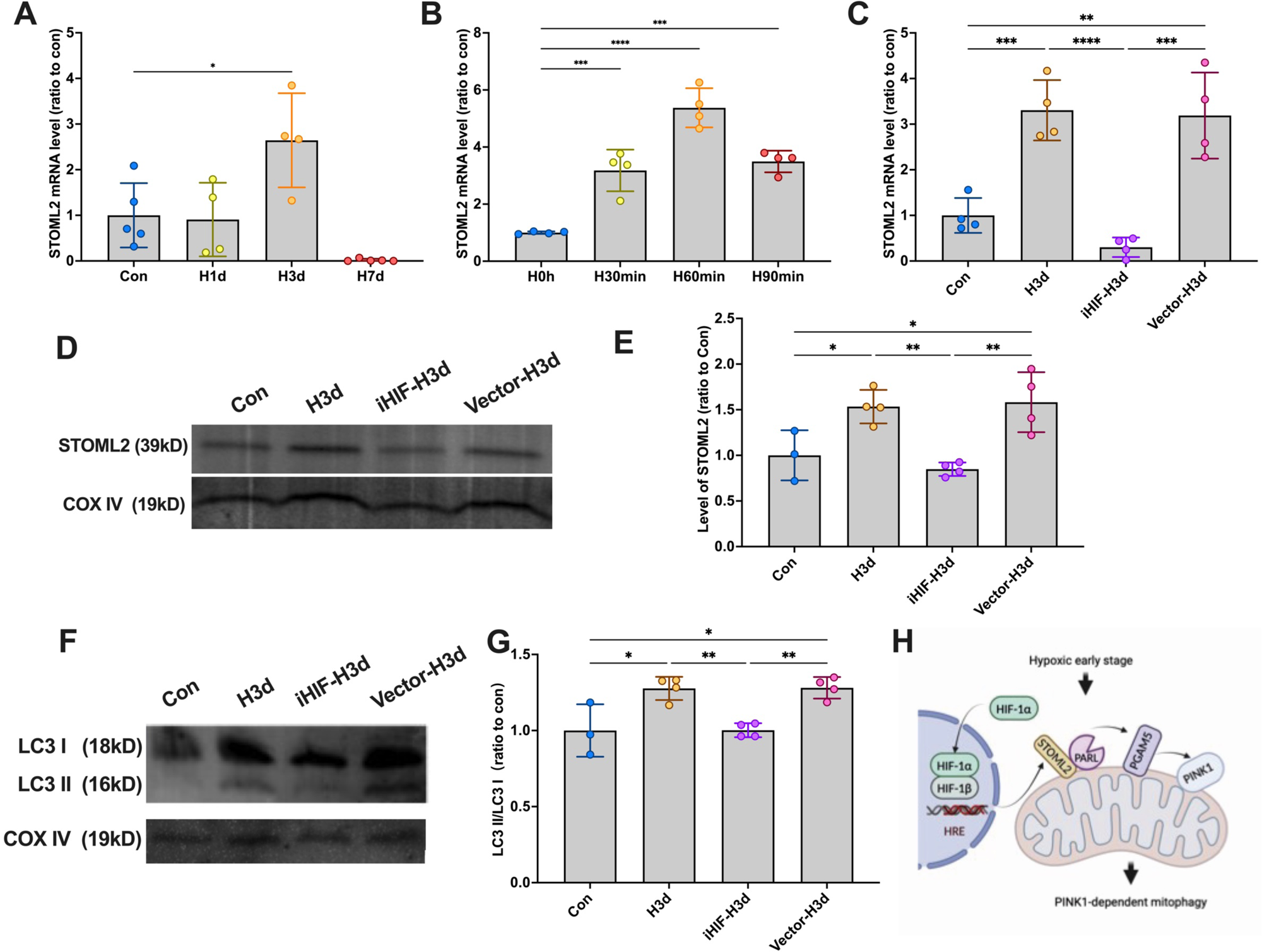
STOML2/PGAM5/PINK1 pathway depended on HIF-1α. **(A)** The levels of STOML2 mRNA were detected by qPCR in mice of each group. **(B)** The levels of STOML2 mRNA were detected by qPCR in cells of each group. **(C-G)** Mice were administered the lentiviral targeting HIF1α and then received continuous hypoxic treatment for 3 days. Mice were grouped in 4 (Con, H3d, iHIF1α-H3d and vector-H3d). **(C)** The levels of STOML2 mRNA were detected by qPCR in mice of each group. **(D)** The levels of STOML2 in mice of each group in the hippocampus mitochondrial protein were detected by western blotting using COX IV as the internal reference. **(E)** Statistical analysis of STOML2 in mice of each group. **(F)** The levels of LC3 in the hippocampus mitochondrial protein were detected by western blotting using COX IV as the internal reference. **(G)** Statistical analysis of LC3 II/I in mice of each group. **(H)** The mechanism diagram of the pathway. Data was analyzed via one-way ANOVA and Tukey’s post-test. *p < 0.05, **p < 0.01, ***p < 0.001, ****p < 0.0001.

Western blot further verified successful HIF-1α knockdown, as nuclear HIF-1α levels were no longer elevated in iHIF-H3d mice (Figure 6D-E). To determine whether HIF-1α acts upstream of mitophagy, we assessed mitochondrial LC3 levels following HIF-1α depletion. Results showed that HIF-1α knockdown abolished mitophagy activation in early hypoxia (Figure 6F-G).

Collectively, our findings demonstrate that early hypoxia promotes HIF-1α nuclear translocation, which in turn upregulates STOML2 expression. This leads to enhanced L-PGAM5 stability and subsequent activation of PINK1-dependent mitophagy. We propose that the HIF-1α/STOML2/PGAM5/PINK1 axis represents a novel hypoxia-responsive signaling pathway that plays a crucial role in neuronal protection against hypoxic injury (Figure 6H).

### Intermittent hypoxia activates PINK1-dependent mitophagy through the HIF-1α/STOML pathway to resist nerve damage

Previous studies suggest that intermittent hypoxia preconditioning (IH) can mitigate cognitive impairment in mice induced by chronic hypoxia. To investigate the underlying protective mechanisms, we examined whether mitophagy contributes to this effect. To assess mitophagy activation, we isolated mitochondrial and cytoplasmic proteins and performed western blot analysis. Results showed a significant increase in LC3-II levels in mitochondria (Figure 7A-B), whereas no such increase was observed in the cytoplasmic fraction (Figure 7C-D). This indicates that IH triggers mitophagy through the intrinsic pathway during early hypoxia.

**Figure 7.**
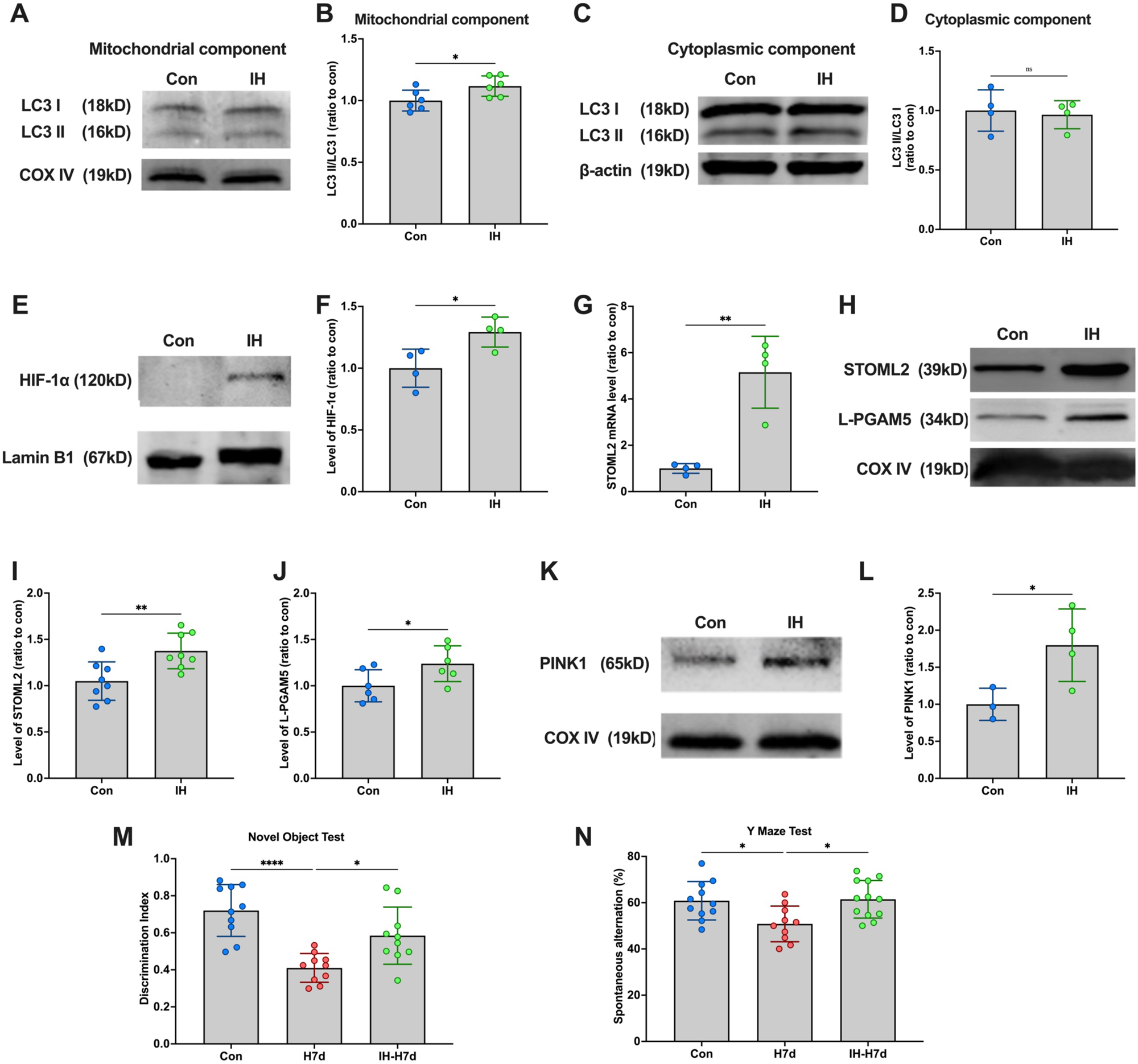
Intermittent hypoxic treatment activates PINK1 dependent mitophagy via HIF-1α/STOML2 pathway. **(A)** The levels of LC3 in the hippocampus mitochondrial protein were detected by western blotting using COX IV as the internal reference. **(B)** Statistical analysis of LC3 II/I in mice of each group. **(C)** The levels of LC3 in the hippocampus cytoplasmic protein were detected by western blotting using β-actin as the internal reference. **(G)** Statistical analysis of LC3 II/I in mice of each group. **(E)** the levels of HIF1α in the hippocampus nucleus protein were detected by western blotting using LaminB1 as the internal reference. **(F)** Statistical analysis of HIF1α in mice of each group. **(G)** The levels of STOML2 mRNA were detected by qPCR in mice of each group. **(H)** The levels of STOML2 and L-PGAM5 in mice of each group in the hippocampus mitochondrial protein were detected by western blotting using COX IV as the internal reference. **(I)** Statistical analysis of STOML2 in mice of each group. **(J)** Statistical analysis of L-PGAM5 in mice of each group. **(K)** The levels of PINK1 in mice of each group in the hippocampus mitochondrial protein were detected by western blotting using COX IV as the internal reference. **(L)** Statistical analysis of PINK1 in mice of each group. **(M-N)** Mice received hypoxic treatment for 7 days after receiving intermittent hypoxic treatment and behavioral tests were performed. Mice were grouped into 3 (Con, H7d, IH+H7d). **(M)** Detection and statistical analysis of Novel Object Test in mice of each group. **(N)** Detection and statistical analysis of the Y Maze Test in mice of each group. Data was analyzed via paired two-tailed Student’s *t*-test. *p < 0.05, **p < 0.01, ****p < 0.0001.

To determine whether IH activates the HIF-1α/STOML2/PGAM5/PINK1 pathway, we analyzed HIF-1α levels in nuclear fractions and found a significant increase after IH treatment (Figure 7E-F). Further western blot analysis of mitochondrial proteins revealed elevated STOML2, L-PGAM5, and PINK1 levels after IH (Figure 7G-H, J-L). Additionally, qPCR analysis confirmed a significant increase in STOML2 mRNA following IH (Figure 7I). These findings suggest that IH activates the HIF-1α/STOML2/PGAM5/PINK1 signaling pathway.

To evaluate whether this pathway contributes to cognitive protection, mice underwent behavioral testing after 7 days of persistent hypoxia following IH treatment. Results from the novel object recognition and Y-maze tests demonstrated that IH significantly alleviated cognitive impairment induced by chronic hypoxia (Figure 7M-N). These findings suggest that IH activates the HIF-1α/STOML2/PGAM5/PINK1 pathway, which in turn induces mitophagy and protects against hypoxia-induced cognitive decline. This pathway may serve as a potential neuroprotective mechanism in hypoxic conditions.

## Discussion

Our findings demonstrate that PINK1-dependent mitophagy serves as a critical protective mechanism in early hypoxia, enabling neurons to selectively eliminate damaged mitochondria and maintain cellular homeostasis (Figure 8). This process is initiated by the stabilization of HIF-1α, which upregulates STOML2 expression, leading to the stabilization of PGAM5 and ultimately activating PINK1-mediated mitophagy. Moreover, this study lays the theoretical foundation for exploring IH as a potential clinical intervention to enhance neuroprotection against hypoxic injury.

**Figure 8.**
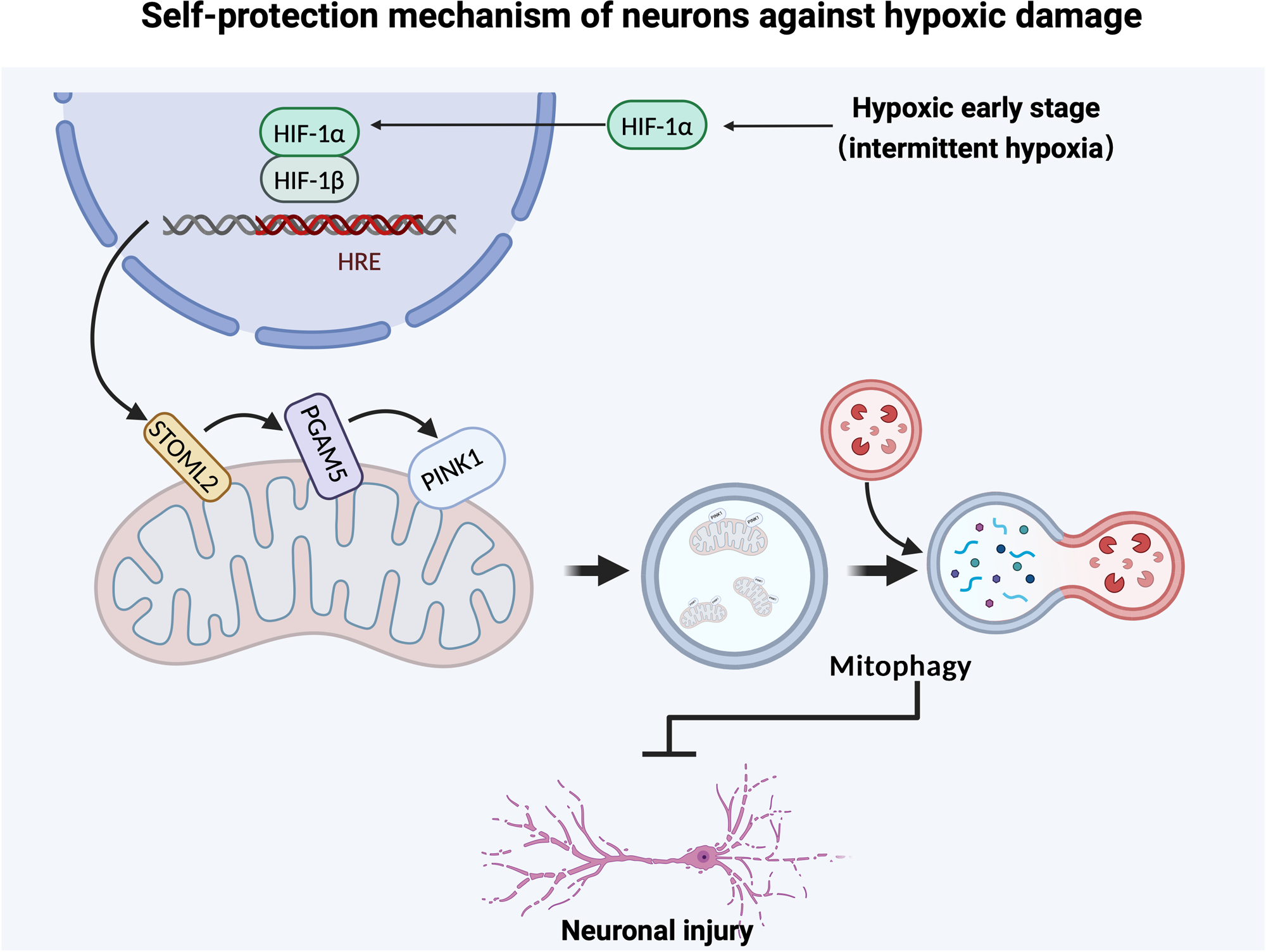
A novel neuronal hypoxia response mechanism. In the early stage of hypoxia or intermittent hypoxia treatment, PINK1-dependent mitophagy is activated by HIF-1α/STOML2-signaling to protect against hypoxia-induced neuronal injury.

During early-stage hypoxia, mitophagy is transiently activated to remove dysfunctional mitochondria, allowing cells to adapt to reduced oxygen levels. However, prolonged hypoxia alters mitochondrial dynamics and cellular stress responses, causing mitophagy to shift from a protective adaptation to a maladaptive process. If mitophagy is sustained for too long, excessive mitochondrial clearance can lead to bioenergetic failure, cellular dysfunction, and even apoptosis. Previous studies have shown that the phosphorylation of FUNDC1 decreases, resulting in diminished mitophagy activity (Tang *et al*, 2024). Similarly, studies in neurons have demonstrated that BNIP3/NIX-mediated mitophagy is transiently activated but suppressed as cells enter hypoxia-induced apoptosis (Chen *et al*, 1999). These findings highlight the need for a tightly regulated mitophagy response, balancing mitochondrial clearance with cellular survival. In this study, we confirm that the HIF-1α/STOML2/PGAM5/PINK1 pathway functions as a novel and effective mechanism for mitophagy activation under hypoxia, as evidenced by knockdown experiments targeting key molecules in the pathway.

Studies have indicated that stabilized HIF-1α enhances mitophagy, reinforcing its role in maintaining mitochondrial homeostasis(Hu *et al*, 2020). Our findings confirm that hypoxia-induced mitophagy is HIF-1α-dependent, underscoring the importance of this regulatory axis. Notably, HIF-1α activation is transient (Randle *et al*, 2024)—it peaks within 4–8 hours and declines after 12–24 hours in HeLa cells (Jewell *et al*, 2001). HIF-1α provides short-term neuroprotection by promoting blood vessel formation (VEGF) and metabolic adaptation(Liu *et al*, 2017), but prolonged HIF-1α activation can lead to blood-brain barrier disruption and edema(Zhang *et al*, 2016). This transient pattern supports our conclusion that mitophagy is similarly short-lived during early hypoxia, with HIF-1α playing a key role in the activation of PINK1-dependent mitophagy.

Given that STOML2 functions as a downstream target of HIF-1α(Zheng *et al*, 2021), we propose a model in which early hypoxia stabilizes HIF-1α, upregulating STOML2 expression, which in turn maintains PGAM5 stability and activates PINK1-dependent mitophagy. Without PGAM5 stabilization, PINK1 is rapidly degraded, leading to impaired mitophagy and exacerbated neuronal injury under hypoxia(Lazarou *et al*, 2015). Interestingly, previous studies have also shown that PGAM5 regulates mitochondrial fission by modulating DRP1 phosphorylation(Pedrera *et al*, 2025) , indicating that its function in mitophagy may extend beyond PINK1 stabilization. Future research should explore whether PGAM5 influences mitochondrial dynamics, particularly the balance between mitophagy and mitochondrial fission-fusion regulation in response to hypoxic stress.

FUNDC1 is the canonical pathway that increases mitophagy after hypoxia. However, in our model, this process relies on PINK1. Unlike FUNDC1, which directly interacts with LC3(Qin *et al*, 2022), PINK1 recruits Parkin, an E3 ubiquitin ligase, which ubiquitinates mitochondrial proteins to initiate mitophagy (Ling *et al*, 2025). PGAM5 is involved in both pathways but plays distinct roles. PGAM5 functions as a phosphatase, dephosphorylating FUNDC1 to enhance its interaction with LC3 under hypoxia (Li *et al*, 2024). In contrast, PGAM5 stabilizes PINK1, preventing its degradation and facilitating mitophagy activation. Both pathways are crucial for maintaining mitochondrial homeostasis during hypoxia but operate through different mechanisms. Additionally, PINK1-dependent mitophagy can be triggered by other stressors, such as IH preconditioning.

In this study, we observed that after IH preconditioning, no cognitive decline was observed in mice following 7 days of continuous hypoxia. This neuroprotective effect was associated with the activation of the HIF-1α/STOML2/PGAM5 pathway, which subsequently enhanced PINK1-dependent mitophagy, promoting mitochondrial quality control and neuronal survival. Previous studies have shown that IH confers neuroprotection through multiple mechanisms: In models of cerebral ischemia, IH promotes mitochondrial biogenesis and prevents oxidative stress-induced apoptosis(Su *et al*, 2022); studies in stroke models have demonstrated that IH increases VEGF expression, improving cerebral blood flow and neuronal survival(Peng *et al*, 2024); IH preconditioning suppresses pro-inflammatory cytokines (TNF-α, IL-6) and enhances antioxidant enzyme activity(Wang *et al*, 2025), thereby protecting neurons from hypoxia-induced damage. Notably, this study provides the first evidence demonstrating that IH specifically activates the HIF-1α/STOML2/PGAM5 axis, revealing a novel regulatory mechanism linking IH to enhanced mitophagy and neuroprotection.

In conclusion, we propose that the stabilization of HIF-1α initiates a novel process that enhances STOML2 expression, promotes PGAM5 stability, and ultimately activates PINK1-dependent mitophagy. This pathway represents a potential new target for treating hypoxia-related diseases. Furthermore, IH may mimic the early stages of hypoxia and serve as an exogenous activator of the HIF-1α/STOML2/PGAM5/PINK1 pathway, providing neuroprotection. Our findings also offer experimental support for the clinical application of IH, although further studies are needed to confirm its therapeutic potential.

## Acknowledgments

Not applicable.

## Materials and Methods

### Animals

Adult male C57BL mice were purchased from SPF Biotechnology (Beijing, China). All animals were housed at room temperature under a 12/12 h light/dark cycle and had free access to food and water. All animal experiments were approved by the Animal Care and Use Committee of the Institute of Animal Management, Capital Medical University (permit no. AEEI-2022-073), and conducted in accordance with ethical requirements and ARRIVE guidelines.

### Hypoxic treatment

All mice were randomly assigned to the control group and each model group. Hypoxic mice were administered hypoxic treatment in a closed hypoxic chamber (China Innovation Instrument Co., Ltd, Ningbo, Zhejiang, China), which accurately set the desired hypoxic concentration and pattern. For chronic hypoxia, mice were treated continuously with 13% O2 for 1, 3, and 7 days. The hypoxic chamber was opened briefly for food and water additions every 3 days. Intermittent hypoxic mice were treated with 10 cycles of 5-min 13% O2 (hypoxia) and 5-min 21% O2 (normoxia) per day for 14 days. The IH-H7d group was followed by an additional continuous 3 days hypoxic treatment after IH treatment.

### Lentivirus Treatment

To investigate the knockdown effects of four target molecules (HIF1α, STOML2, PGAM5, and PINK1), along with a negative control (vector), five lentivirus treatment groups were established, with 10 animals per group: H3d-vector group: Experimental animals were injected with lentivirus via stereotactic injection and subjected to continuous hypoxia (13% O₂) for 3 days. H3d-iHIF group: Lentivirus was administered via stereotactic injection to specifically knock down HIF1α, followed by 3 days of continuous hypoxia (13% O₂). H3d-iSTOML2 group: Lentivirus was injected to selectively knock down STOML2, followed by 3 days of continuous hypoxia (13% O₂). H3d-iPINK1 group: Lentivirus was injected to specifically knock down PINK1, followed by 3 days of continuous hypoxia (13% O₂).

### Behavioral tests

The cognitive function of mice in each group was assessed using novel object recognition and Y-maze tests. For the novel object recognition test, a 40 cm × 40 cm × 40 cm lidless rectangular box was used, with a camera positioned overhead. The experiment consisted of three phases: adaptation, familiarity, and testing. In the adaptation phase, each mouse was placed in the apparatus and allowed to explore freely for 5 minutes to acclimate. In the familiarity phase, two identical objects A (old object) were introduced, and mice were given 5 minutes to explore. In the testing phase, one object A was replaced with a novel object (differing in color and shape), and mice were allowed another 5 minutes of exploration. The time spent interacting with both objects was recorded. The discrimination index was calculated as (Time exploring new object − Time exploring old object) / (Time exploring new + old objects). Mice with baseline cognitive impairments were excluded from behavioral tests. All video recordings were analyzed blindly by researchers not involved in conducting the experiments.

The Y maze is typically made of opaque material and shaped like the letter “Y,” consisting of three equal-length arms (usually at a 120° angle). Each arm is approximately 30–40 cm long, 8–10 cm wide, and 15 cm high to prevent animals from escaping. The apparatus is placed in a disturbance-free laboratory environment, and a video tracking system is used to record animal behavior. Before the experiment, animals may be allowed to acclimate to the laboratory environment for about 30 minutes to reduce anxiety. The test begins by placing the animal in the start arm of the Y maze (typically a fixed arm), allowing it to explore freely for a set period (usually 5–10 minutes). During the experiment, the sequence and number of entries into each arm are recorded to calculate the alternation rate. A successful alternation is defined as consecutive entries into three different arms, such as A→B→C. If the animal revisits any arm within three consecutive choices, it is not counted as a successful alternation. The spontaneous alternation rate is calculated as: Spontaneous alternation rate = Number of successful alternations / Total exploration attempts.

### Nissl staining

Brain tissue from the mice were cut into 10 μm sections on a frozen slicer and pasted on a slide. The samples were fixed in 70% ethanol, then sequentially dehydrated in 100%, 90%, 80%, and 70% ethanol for 2 minutes each. After clearing with xylene, the sections were incubated in 1% tar purple (Solarbio, G1430) for 30 minutes. They were then rinsed with distilled water and differentiated in 70% alcohol for several minutes. Dehydration was repeated with 70%, 80%, and 95% ethanol for 2 minutes each, followed by 100% ethanol, before being sealed with neutral gum.

### Western blots

Mouse hippocampus protein lysates were resolved by sodium dodecyl sulfate-polyacrylamide gel electrophoresis (SDS-PAGE) and subsequently immunoblotted onto polyvinylidene difluoride (PVDF) membranes. Membranes were blocked with 5% nonfat milk at room temperature for 1 h. After TBST washing (three times, 5 min per wash), the membranes were incubated with the indicated primary antibodies at 4 °C overnight with shaking. The primary antibodies included: COX IV (Proteintech, 23274-1-AP), LaminB1 (Proteintech, 80906-1-RR), HIF-1α (Abcam, ab228649), STOML2 (Proteintech, 60052-1-AP), PGAM5 (Proteintech, 28445-1-AP), PINK1 (Abconal, A11435), LC3 (Sigma,L7543).After incubation, the membranes were washed three times and then incubated at room temperature for 1 h with secondary antibodies, including IRDye 680RD goat anti-mouse IgG (H + L) (Licor, 926-68070), IRDye 680RD goat anti-rabbit IgG (H + L) (Licor, 926-68071), IRDye 800CW goat anti-mouse IgG (H + L) (Licor, 926-32210), IRDye 800CW goat anti-rabbit IgG (H + L) (Licor, 926-32211). PVDF were scanned using a detection system (Odyssey, USA), and band intensities were normalized to Lamin B1 or COX IV. Statistical analyses were performed using ImageJ and GraphPad software.

### RT-PCR

An RNeasy kit (Qiagen, 74104) was used to extract total RNA from mice hippocampal tissue, and then the Transcriptor High Fidelity cDNA synthesis kit (Roche, 5081963001) was used to reverse transcribe the RNA into cDNA. All operations were according to the instructions. The following primers were used: Stoml2 for: TACAAGGCAAGTTACGGTGTGG; Stoml2 rev: GAGAATGCGCTGACATACTGCT; 18S sense: GTAACCCGTTGAACCCCATT; 18S anti: CCATCCAATCGGTAGTAGCG.

### Cytotoxicity detection

Cytotoxicity was detected by the LDH assay (Roche, 4744926001). The powder was dissolved in ddH2O and mixed thoroughly to make the catalytic solution. Then, 250 μL of the catalytic solution was added to the staining solution (11.25 mL) and mixed thoroughly. Then, 100 μL cell supernatant of each group was added to the new 96-well plate. The LDH reaction solution (100 μL) was added with subsequent incubation at room temperature away from light for 30 min. After the incubation, 50 μL stop solution was added to each well and gently mixed for 10 min. The OD value of each well was measured at 490 nm by a microplate reader.

### Cell Proliferation Assay

Cell proliferation was detected by the CCK8 assay(), including followed steps: (1) Standard Curve: Count the cells in the suspension and prepare a cell concentration gradient. Dilute with culture medium to create 4-7 gradients, incubate overnight, then remove the medium and add fresh medium with CCK-8 reagent. After 1 hour, measure OD to create a standard curve. (2) Cell Seeding: Seed cells at the optimal density determined, and incubate overnight. (3) Add CCK-8: Remove the original medium, and add 100 μl of medium with CCK-8. (4) Incubation: Incubate for 1 hour, then transfer the hypoxia group to a 1% O2 incubator. (5) OD Measurement: Measure OD at 450 nm using a microplate reader.

### Cell death detection

PI/Hoechst detection was used to detect the cell death rate. Hoechst labels all cells as blue fluorescence, and PI labels only dead cells as red fluorescence. Therefore, the ratio of red to blue can be used to calculate the cell death rate. After the cells were treated, the original medium was discarded, and the cells were rinsed three times with PBS. The PI (Sigma, P4170) and Hoechst (Sigma, B2261) mixture was added into the cell culture well and incubated at 37 °C for 10 min under dark conditions. The cells were removed from the incubator, the mixture of PI and Hoechst was discarded, and the cells were rinsed three times with PBS. Confocal microscopy was used for observation and imaging.

### MitoTracker Staining and Flow Cytometry Detection

Using MitoTracker Green probe kit purchased from Thermo Corporation. (1) Prepare staining solution: dilute 1mM MitoTracker stock in serum-free medium to a working concentration of 150nM. (2) Staining: Once cells reach the desired density, discard the old medium, add pre-warmed MitoTracker solution, and incubate for 10 minutes in the dark. (3) Remove staining solution: Replace with regular medium. (4) Flow cytometry: Digest, centrifuge, and collect cells to create a single-cell suspension. Perform fluorescence detection using a flow cytometer with the FITC channel.

### Mitophagy Inhibition

(1) Preparation of Mdivi-1 Working Solution: Dissolve Mdivi-1 (purchased from Selleck, S7162) in DMSO to prepare a stock solution and store it at -20°C in the dark. Before use, dilute the stock solution in complete culture medium to a final concentration of 10 µM. (2) Cell Culture: Seed cells in a 96-well plate and incubate at 37°C with 5% CO₂ until they reach the appropriate density. (3) Cell Treatment: Remove the original culture medium and replace it with fresh medium containing Mdivi-1 at a ratio of 10 µl per 1 ml of medium. Pre-treat the cells for 30 minutes. (4) Subsequent Assays: Perform cell viability and functional assays such as CCK-8.

### Statistical analysis

Excel and GraphPad Prism 9.0 software were used for data preservation, recording, statistics, and analyses. Image data were analyzed by ImageJ and other software. All results were analyzed using the t-test, one-way ANOVA, and two-way ANOVA as appropriate. Data are expressed as the mean ± standard error (mean ± SEM), with P ≤ 0.05 as a significant difference. In animal behavioral tests, n ≥ 10; in protein detection experiments, such as western blot and immunofluorescence, n ≥ 3.

## Abbreviations

CNS: central nervous system
AD: Alzheimer’s disease
PD: Parkinson’s disease
ROS: reactive oxygen species
PINK1: PTEN-induced kinase 1
OMM: outer mitochondrial membrane
FUNDC1: FUN14 domain-containing 1
I/R: ischemia/reperfusion
IH: intermittent hypoxia
Drp1: dynamin-related protein 1
HIF-1α: hypoxia inducible factor-1α;

## Declaration of Artificial Intelligence (AI)

During the writing process of this article, we used ChatGPT-4o for language refinement and optimization to enhance readability and fluency. However, all research content, data analysis, and conclusions were independently conducted by the authors. ChatGPT-4o was solely used for language enhancement and did not influence the scientific integrity or authenticity of the study.

---

**Figure S1. PINK1 overexpression enhances the level of mitophagy. (A)** The levels of PINK1 in mice of each group in the hippocampus mitochondrial protein were detected by western blotting using COX IV as the internal reference. **(B)** Statistical analysis of PINK1 in mice of each group. **(C)** The levels of LC3 in the hippocampus mitochondrial protein were detected by western blotting using COX IV as the internal reference. **(D)** Statistical analysis of LC3 II/I in mice of each group. Data was analyzed via one-way ANOVA and Tukey’s post-test. *p < 0.05, **p < 0.01, ***p < 0.001, ****p < 0.0001.

**Figure S2. STOML2 acts as an upstream regulator of PGAM5. (A)** The levels of STOML2 in mice of each group in the hippocampus mitochondrial protein were detected by western blotting using COX IV as the internal reference. **(B)** Statistical analysis of PINK1 in mice of each group. Data was analyzed via one-way ANOVA and Tukey’s post-test. *p < 0.05.

